# Design and Validation of a Wearable Imaging System for Automated Wound Monitoring in Porcine Model

**DOI:** 10.1101/2024.12.18.629174

**Authors:** Wan Shen Hee, Maryam Tebyani, Prabhat Baniya, Celeste Franco, Gordon Keller, Fan Lu, Houpu Li, Narges Asefifeyzabadi, Hsin-ya Yang, Guillermo Villa-Martinez, Kaelan Schorger, Koushik Devarajan, Alexie Barbee, Cristian Hernandez, Tiffany Nguyen, Marcella Gomez, Roslyn Rivkah Isseroff, Marco Rolandi, Mircea Teodorescu

**Affiliations:** Department of Electrical and Computer Engineering, University of California Santa Cruz; Department of Dermatology, School of Medicine, University of California Davis; Department of Ophthalmology & Vision Science, University of California Davis; Department of Applied Mathematics, University of California Santa Cruz; Department of Biomolecular Engineering, University of California Santa Cruz; Genomics Institute, University of California Santa Cruz

**Keywords:** wearable imaging device, wound monitoring, wound analysis, wound stage prediction, porcine model

## Abstract

Effective wound monitoring has the potential to guide treatment regiments and improve healing outcomes, yet current clinical assessment methods remain largely subjective and labor-intensive. To address this challenge, we present a high-resolution wearable imaging system designed for continuous wound monitoring. The system integrates a 64 MP camera with a plano-convex lens housed in an enclosure measuring 73 mm in diameter and 36.1 mm in height, and features a custom printed circuit board (PCB) for programmable LED illumination. The 3D-printed device enclosure is designed to accommodate a silicone bioelectronic device and can be securely attached using a commercially available ostomy skin barrier. In porcine model validation studies, the system successfully captured daily wound progression over periods up to 7 days. The captured images were wirelessly transmitted to a processing unit where DeepMapper, a machine learning algorithm, processed z-stacked images and performed multi-level feature extraction to predict wound healing stages and indicate potential complications such as infection. This imaging system enables automated analysis of wound progression and supports the development of smart wound care platforms for personalized treatment strategies. The integrated design approach demonstrates the feasibility of creating compact, high-resolution imaging systems suitable for clinical wound monitoring applications.

## 1 Introduction

The skin serves as the human body’s primary defense against the external environment, including pathogens, chemicals, and extreme temperature fluctuations. When this barrier is compromised through injury, the resulting wounds can range in severity from superficial abrasions to life-threatening conditions.^1, 2^ The clinical and economic impact of wound care is substantial - in the United States alone, more than 8 million people require professional wound care annually, contributing to healthcare costs exceeding $28 billion.^3, 4^ Efficient wound healing and monitoring is crucial in reducing hospitalizations, lowering overall healthcare expenditures, and improving patient outcomes.

Extensive research has been conducted to classify and characterize the complex process of wound healing. Wounds are typically understood to progress through four overlapping stages of healing: hemostasis, inflammation, proliferation, and maturation.^2, 5–8^ Wounds are classified as acute or chronic based on their healing trajectory and underlying causes.^1, 2, 8, 9^ Acute wounds progress sequentially through the four stages without significant delay, leading to full recovery. In contrast, a chronic wound arises when one or more stages of healing are disrupted, requiring extended professional treatment and specialized care.^1, 6, 8^

Despite the importance of wound healing, current clinical assessment primarily relies on physicians’ expertise, which is inherently labor-intensive and subjective. This dependence on visual inspection and clinical experience can result in delays in detecting complications and initiating timely interventions. Although various diagnostic tools have been developed to assist in wound assessment, these tools have yet to reliably predict healing stages, leaving clinicians to rely heavily on their judgment for accurate diagnosis.^10^

Recent advances in biomedical imaging and wearable technologies offer promising solutions to these challenges in wound management.^8^ Wearable devices enable continuous, minimally invasive to non-invasive monitoring, providing real-time, high-resolution insight into wound healing. These systems allow clinicians to track wound progression in detail, facilitating early detection of complications and enabling timely interventions when healing deviates from the expected trajectory.^11–13^ By combining wearable devices with machine learning algorithms, these devices can automate the analysis of wound features,^14^ predict healing stages^15^ and healing time,^16^ create personalized precision treatment plans^17^ and ultimately reduce wound healing time.^18^ These technologies extend specialized wound care beyond traditional healthcare settings, reducing the frequency of clinical visits and associated costs through remote patient monitoring, particularly benefiting patients in remote areas or those with limited mobility.^19^ Further, bioelectronic actuators enable the programmable delivery of personalized treatments, such as charged biomolecules,^13^ to complement monitoring efforts. The integration of wearable technologies and artificial intelligence presents a promising approach toward more precise, personalized, and accessible wound care.

In this paper, we present a high-resolution wearable imaging system developed for continuous wound monitoring. The camera system has been designed for use with a machine learning algorithm and a silicone-based bioelectronic actuator to provide real-time healing assessment and guide wound treatment. The system captures 32MP images of the wound area at regular intervals, allowing for detailed tracking of changes in wound size, color, and texture. These images are analyzed using DeepMapper, an attention-based autoencoder proposed by Lu et al.^20^ to predict the wound stage. We begin by describing the system design and architecture, including the device and controller, along with the calibration of the optical system. We evaluate the device performance by characterizing the resolution and imaging subdermal porcine tissue *ex-vivo*. Next, we detail the device operation protocol for multi-day *in-vivo* experiments and describe the use of DeepMapper for wound stage prediction. Finally, we present results from porcine wound experiments, demonstrating the system’s ability to continuously monitor wound progression and detect complications throughout the healing process.

## 2 System Design

The proposed system, shown in Figure 1, consists of a custom-designed imaging system enclosed together with a polydimethylsiloxane (PDMS) based bioelectronic actuator. The controller PCB uses WiFi to connect to an off-board processing unit that manages device operations, including programmed image capture, and receives locally stored wound images. Once the images are received, a machine learning algorithm on the processing unit uses them to generate a wound stage prediction for the corresponding wound. The device and controller are powered by a power bank housed in a pouch, as shown in Figure 4b.

**Fig 1.**
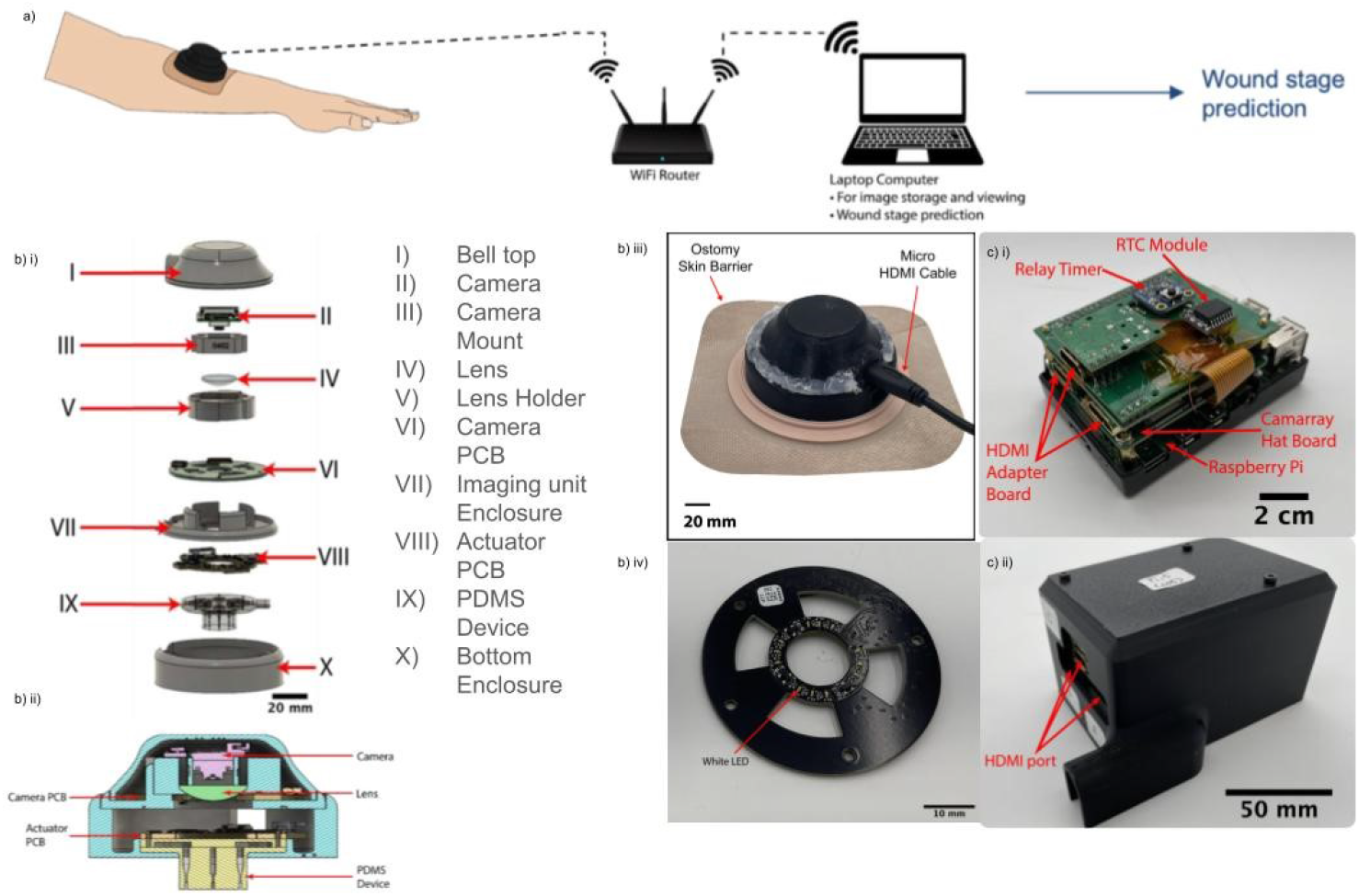
a) System overview; b)(i) Exploded view, (ii) Cross section of the device, (iii) The assembled device attached to an ostomy skin barrier, (iv) The bottom of the camera PCB; c) (i) The controller stack and (ii) controller enclosure.

### 2.1 Device and Controller Design

The wearable imaging system is enclosed together with the bioelectronic actuator. The actuator’s operating principle has previously been used to deliver an electric field and charged biomolecules *in-vitro*^21^ and for expedited wound healing in murine models *in-vivo*.^13, 22^ Figure 1b(i) and Figure 1b(ii) show exploded and cross-sectional views of the device’s CAD design. The device integrates optical and electronic components in a compact design that includes a camera, a plano-convex lens, a custom camera printed circuit board (PCB), and the PDMS-based bioelectronic actuator. The camera PCB features 12 LED banks for illumination and interfaces from the camera’s 15-pin flexible flat cable (FFC) connector to a micro-HDMI output for robust connection to the controller. These components are housed within 3D-printed enclosures (designed in Autodesk Fusion 360^23^). Figure 1b(iii) shows the device fixed to an ostomy skin barrier, which secures it to the wound during *in-vivo* experiments. The device has a circular footprint of 73 mm in diameter and stands 36.1 mm above the skin surface.

To provide magnification and high resolution for wound imaging, the system uses a plano-convex lens in combination with an Arducam 64 MP Hawkeye camera. In the proposed design, the camera and LEDs interface with the controller via the custom PCB, which incorporates a micro-High Definition Multimedia Interface (HDMI) connector, a 15-pin FFC connector, and illumination LEDs. Previous iterations of the design, shown in the supplementary materials, used two separate breakout boards: one for the micro-HDMI connector and another for the illumination LEDs via a 6-pin flexible printed cable (FPC). The current design consolidates these breakout boards onto the custom PCB, reducing the number of components and minimizing the device’s overall footprint which, in turn, improves reliability and lowers the risk of mechanical failure. Figure1b(iv) shows an isometric bottom view of the custom camera PCB, highlighting the white LEDs and the associated electronics used for their control. Further details on the design can be found in the supplementary materials.

The Figure 1c(i) shows the controller, which consists of a three-tiered architecture. At the base, a Raspberry Pi 4b serves as the core microcontroller unit which powers and controls all connected electronics. On top of the Raspberry Pi, a CamArray HAT multiplexing board is mounted, which allows control of up to four camera systems; we have connected up to three in our testing. Above, a custom PCB, named the Pi Shield, houses additional electronics that expand the Raspberry Pi’s functionality. The Pi Shield includes three HDMI breakout boards, which convert the flexible flat cable (FFC) from the camera into a more robust HDMI connection. This connector significantly improves the mechanical integrity of the system, as FFCs are prone to disconnection under stress. The HDMI breakout boards also allow the Raspberry Pi’s General Purpose Input/Output (GPIO) pins to directly control the LEDs on the camera PCB. In addition, the Pi Shield contains a real-time clock (RTC) module to provide accurate timestamps for the captured images and a relay timer to wake the Raspberry Pi at preset intervals for wound imaging.

The controller is encased in a 3D-printed housing with access ports for the micro-HDMI connectors during experimental procedures, as shown in Figure 1c(ii). Further information on the controller design can be found in the supplementary materials.

### 2.2 Device Validation

Figure 2 details the device design and validation process. Figure 2a presents the proposed configuration of the optical components within the device. These components include the Arducam 64 MP Hawkeye camera, responsible for capturing high-resolution wound images; a plano-convex lens for image magnification; an LED ring to illuminate the image target; and a mock PDMS to represent the bioelectronic device. All components are aligned along a common optical axis. This configuration is also used as a calibration set-up.

**Fig 2.**
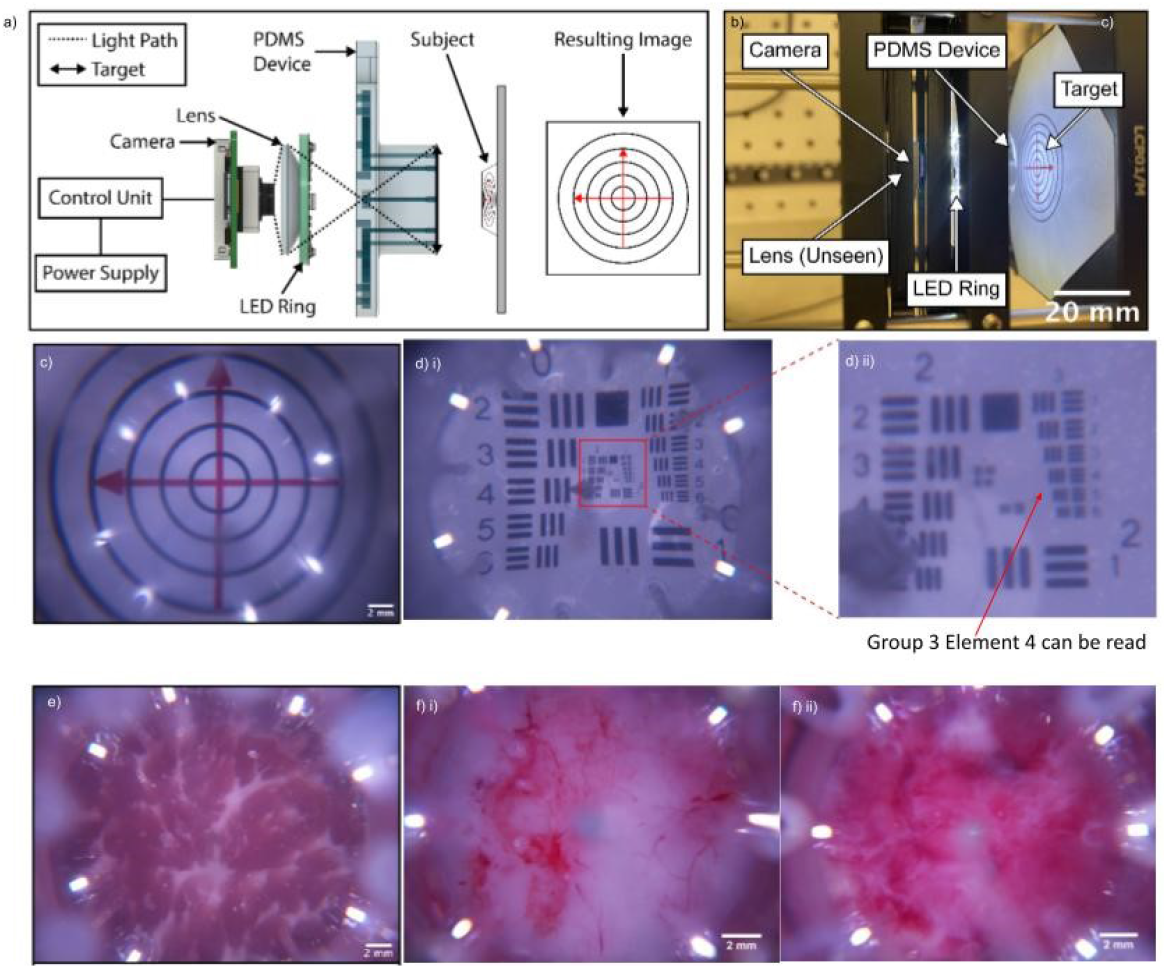
Imaging system design and calibration. a) Schematic diagram of the optical system; b) Calibration setup to determine the placement of each optical component and the focal range of the device; c) Test target image from calibration setup, showing the optical field of view using perpendicular arrows of 20 mm length; d) Resolution characterization using USAF 1951 target (i) Full target image, (ii) Magnified view showing smallest resolvable line pairs (Group 3 Element 4); e) *Ex-vivo* validation using the assembled device to image porcine subdermal tissue; f) *In-vivo* porcine wound monitoring using the assembled device. (i) Day 0 wound bed showing initial vascular response post-wound generation, (ii) Day 6 wound bed showing tissue changes and fluid accumulation.

To determine the optimal positioning of these optical elements, the components were mounted on an optical stage as shown in Figure 2a. Images of a test target were captured using only the optical components on the stage. The test target is color-printed and consists of two perpendicular arrows, each 20 mm in length, intersecting at the center of several concentric circles with radii increasing in 5 mm increments, as shown in Figure 2a. This design was chosen to match the expected in vivo wounds, which have a 20mm diameter.

To achieve a compact design without compromising image quality, images of the test target were taken at various object and image distance combinations to identify the optimal positioning of each component. Figure 2c shows an image of the test target captured during the calibration, where the calibration produces an in-focus image with a 20 mm diameter.

After determining the positioning of the optical components, the device enclosure was designed and 3D-printed. The optical components were then assembled into the enclosure to form the complete device. The assembled device was evaluated using a USAF 1951 resolution target, shown in Figure 2d(i). The device successfully resolved the line pairs up to Group 3, Element 4, as shown in Figure 2d(ii). This element corresponds to a spatial frequency of 11.31 line pairs per millimeter (lp/mm), indicating a resolution of 44.19 *µm* for the device.

*Ex-vivo* testing was then conducted by imaging a piece of porcine sub-dermal tissue to demonstrate the device’s capability for wound imaging in porcine models. The results, presented in Figure 2e show the muscle and fat layers in the porcine sub-dermal tissue. Finally, Figure 2f shows images of a porcine wound captured under *in-vivo* conditions during a 7-day experiment. Figure 2f(i) represents the wound at the start of the experiment (Day 0), while Figure 2f(ii) shows the wound at the end of the experiment (Day 6).

### 2.3 Device Operation

The high-level device communication between the deployed system and remote computer is illustrated in Figure 3a. A single controller unit is used to control up to three devices simultaneously and communicate wirelessly with a computer through a local WiFi network, set up using a commercial router. This configuration allows for flexible customization of camera and electronic operations through programming. During *in-vivo* experiments, two controllers were deployed and individually powered by a dedicated battery pack on either side of the porcine model, ensuring balanced weight distribution. The battery packs were replaced daily to ensure sufficient power.

**Fig 3.**
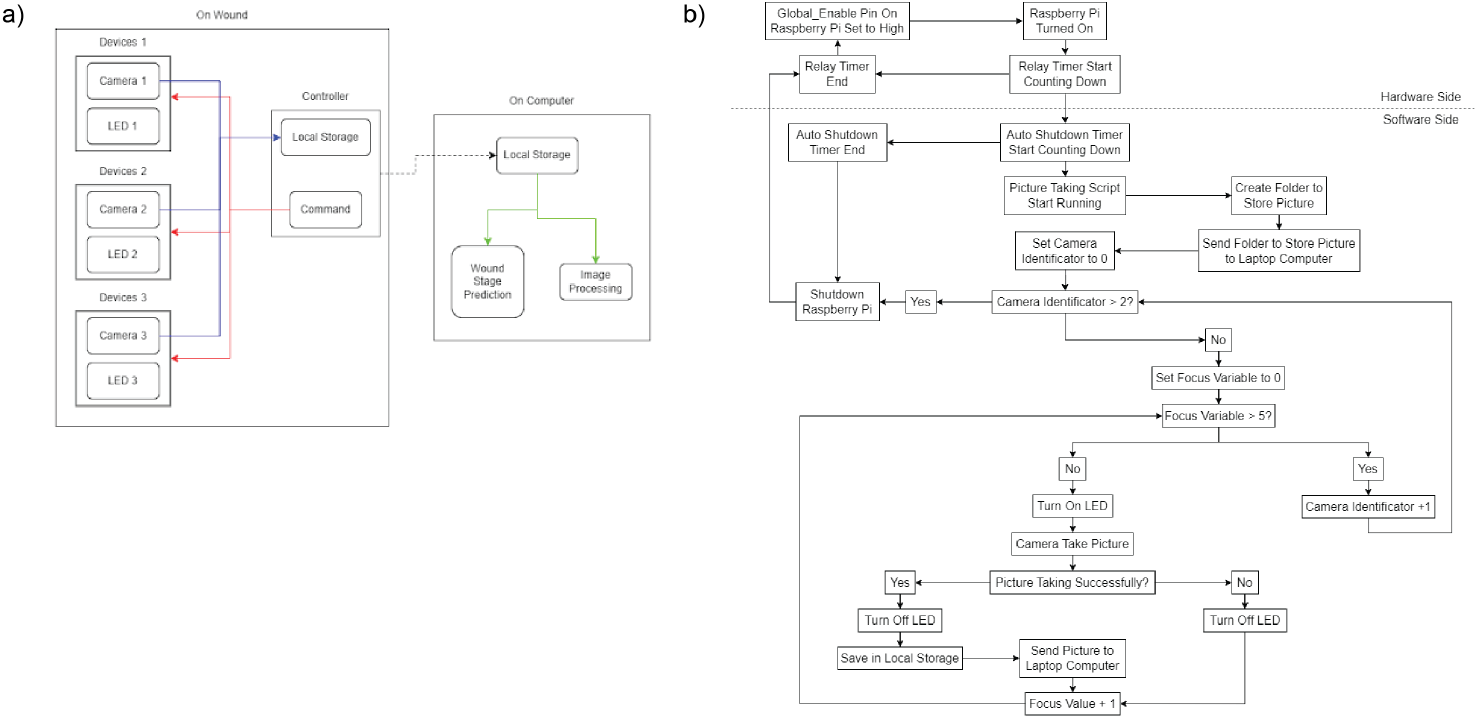
System architecture and operational flowchart. a) High-level device communication showing the interaction between wound-mounted devices (each consisting of a programmable camera and LED) and the remote computer. Solid lines represent wired connections, and the dashed line indicates wireless data transfer from the controller for wound prediction and image processing; b) Detailed operation flowchart of the controller’s hardware and software-based operational sequence, illustrating the automated image acquisition process including device initialization, focus adjustment, image capture, and data storage procedures.

While each controller can control up to three devices, factors such as the pig’s size and the number of physiological control wounds required were used to determine the number of connected devices deployed per controller. Thus, the number of devices per side per pig fluctuates, and the controller’s software is parameterized to allow for this variation.

For image acquisition, we used libcamera,^24^ which natively supports Arducam’s camera systems on the Linux operating system. During the experiments, wound images were captured every two hours. According to,^25^ the first stage of wound healing, which is the hemostasis phase, occurs immediately after the injury and typically concludes within a few hours. Following the completion of the hemostasis phase, the inflammatory phase begins and generally lasts up to 72 hours post-injury. Therefore, the proposed imaging interval provided sufficient imaging cycles each day to capture changing in the wound dynamics while optimizing power consumption. A hardware relay timer on the controller was programmed to send a trigger signal every two hours to boot up the controller. Once the controller completed its startup sequence, the imaging cycle began automatically.

At the start of each imaging cycle, the controller signals the devices to activate their LEDs and initiate image acquisition. Once an image is taken, the LEDs are turned off to conserve power, and the image is stored locally on the controller before being transmitted wirelessly to the computer. The image acquisition system captures a z-stack, which is a series of images taken at multiple focal planes. This is essential for imaging the non-uniform surface topology of wound beds with varying depths and skin thickness. This is enabled by the motorized lens of the Arducam 64 MP Hawkeye camera, which allows programmable adjustment of the focal plane. The system-level image acquisition process proceeds sequentially across devices, with each device capturing a complete six-image z-stack before moving to the next device. Using the z-stack method allows us to process individual images with the best focus in post-processing. Once all devices have completed image acquisition, the imaging cycle concludes with the controller powering down to conserve energy. A new imaging cycle automatically begins every two hours throughout *in-vivo* experiments. Figure 3b illustrates the detailed operational flowchart for the controller and devices.

### 2.4 Wound Stage Prediction

The device captures a set 6 z-stack wound images, and wirelessly transmitted to the remote computer. Then, the z-stack images are merges into a single image, referred to as the “original picture” in Figure 4c. The image was color corrected to the original picture to restore the wound’s true color.^20^ A sample color-corrected image is shown in Figure 4d. The color-corrected picture was down-sampled before it was used for the wound prediction shown in Figure 5b.

**Fig 4.**
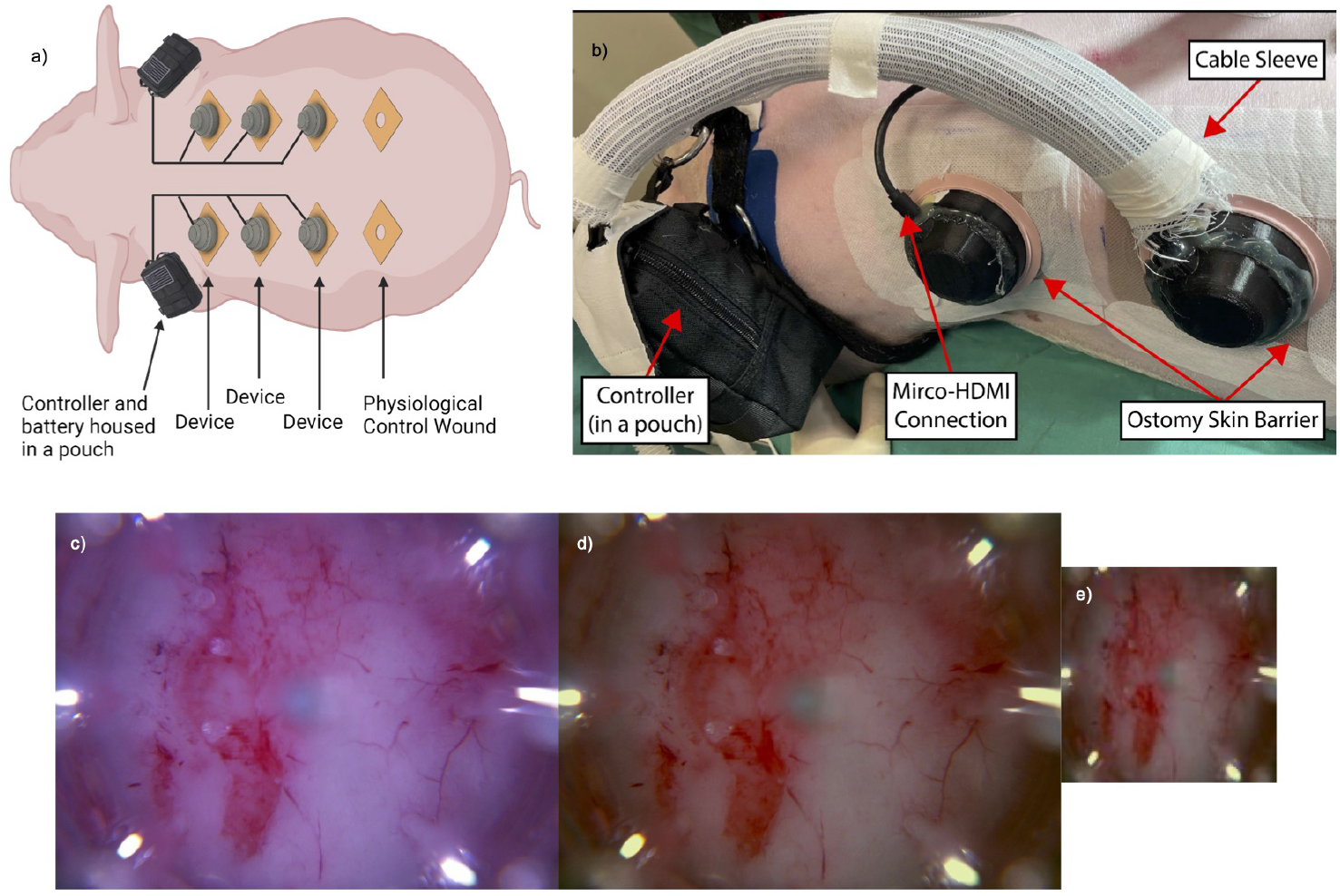
*In-vivo* experimental setup and image processing demonstration. a) Schematic showing device placement on the porcine model, including three active devices and one control wound per controller; b) Photograph of deployed system including the controller housed in a pouch, cable management, and wound interfacing using ostomy skin barriers; (c) Original Day 0 wound image, (d) Same image after color correction processing, and e) Down-sampled version of the wound image used for healing stage prediction in Figure 5.

**Fig 5.**
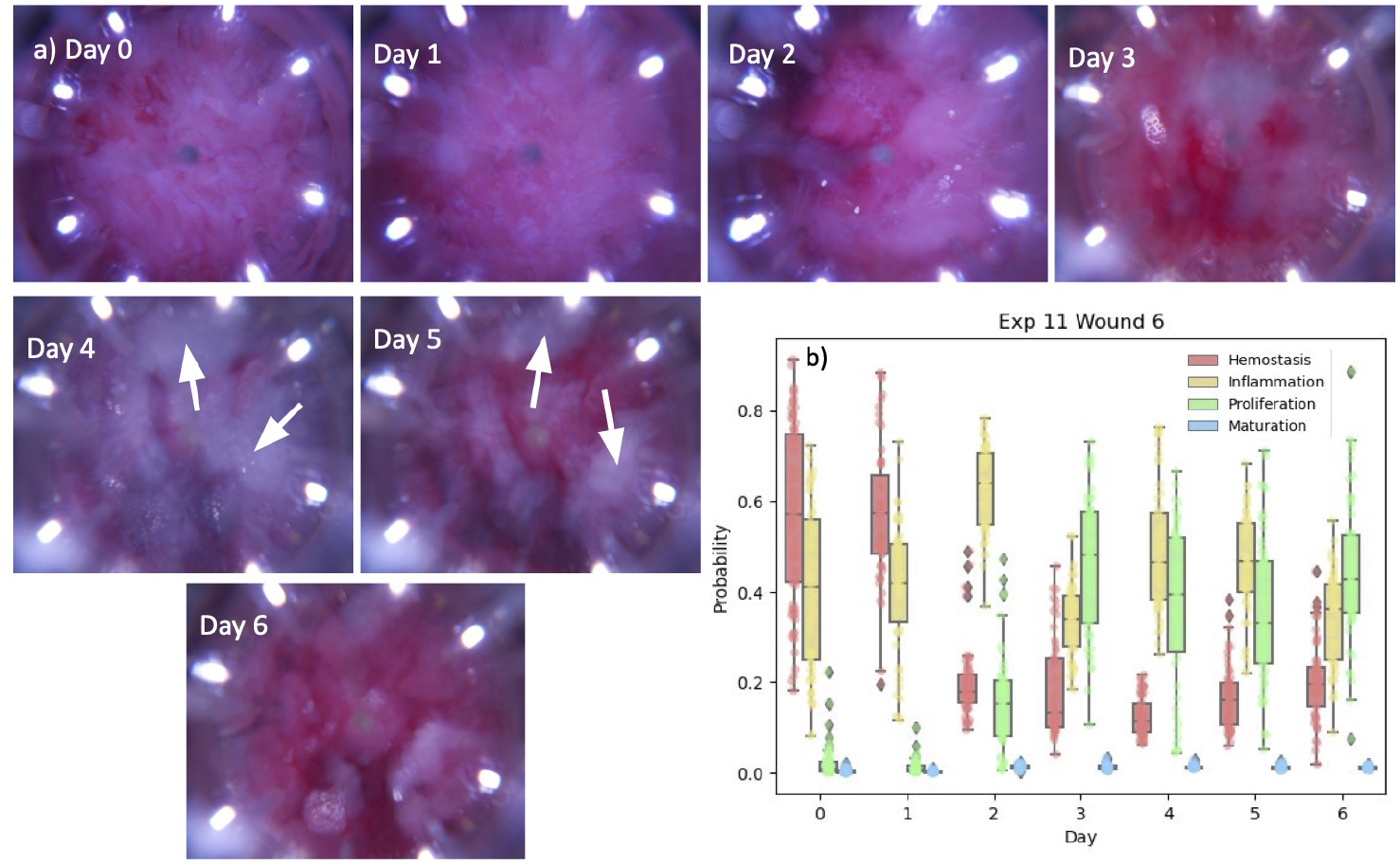
Temporal wound monitoring and wound stage prediction. a) Representative images of a wound captured by the wearable device over a 7-day *in-vivo* experiment. The white arrows indicate potential signs of infection on Days 4 and 5. b) Box plots showing the machine learning model predictions of wound healing stage probabilities over time. Each data point represents the model’s probability output for individual images.

**Fig 6.**
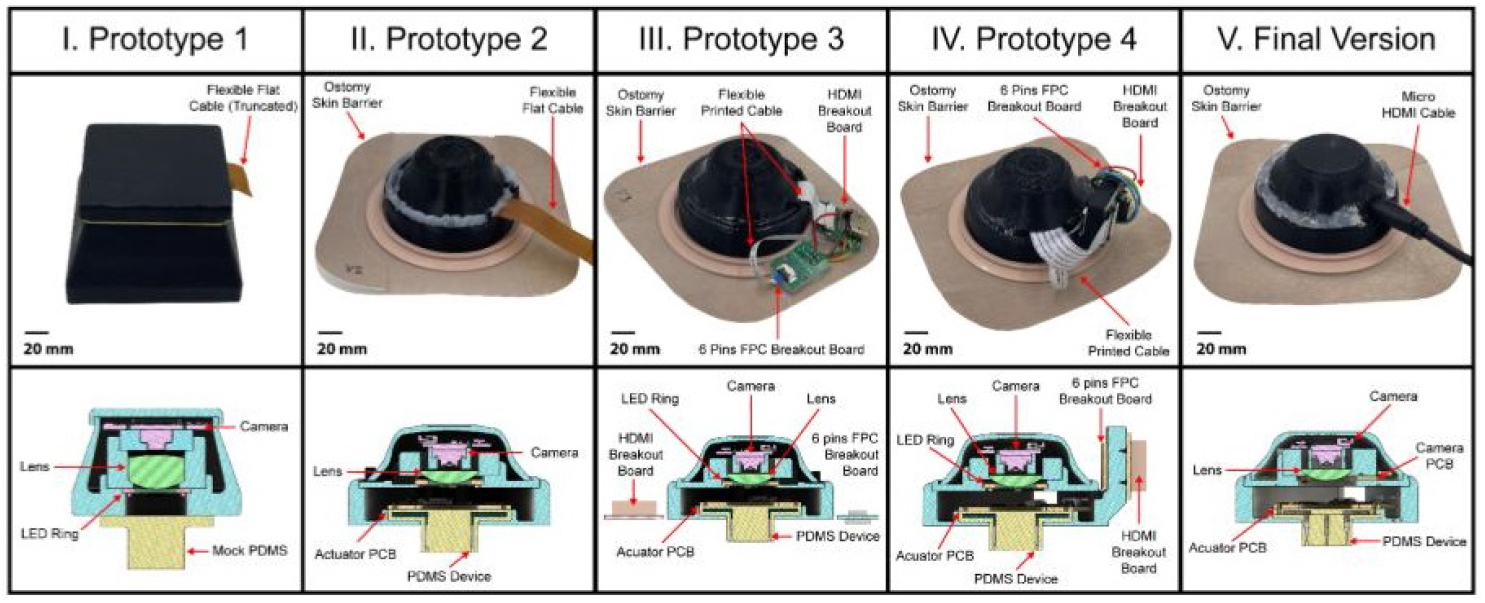
Prototypes and CAD design of the wearable device are shown, with the final version presented in the main text. The iterative design process streamlined production while maintaining the functionality described in the main text.

## 3 Results and Discussion

Figure 5 shows a series of in-vivo images collected by the assembled device from a porcine model 7-day experiment. Figure 5a shows one representative figure per day. These images are captured through the integrated bioelectronic actuator. Although the actuator’s optical clarity is optimized during fabrication, minor occlusions from its features are visible on the image perimeter. Figures 5b shows the wound stage prediction (Hemostasis, Inflammation, Proliferation and Maturation^20^).

Signs of infection became noticeable on Day 4 and Day 5, indicated by the white arrows, which aligns with the higher predicted inflammation values on these days. By Day 6, the visual signs of infection dissipate which is also reflected in the model output prediction. A wound swab taken post-experiment indicated a relatively high colony-forming unit (CFU) count of 1.1×10^3^/mL, although the inflammation predicted on Day 6 is lower. The re-epithelialization rate for this wound was recorded at 27.383%.

## 4 Materials and Methods

We perform *in-vivo* testing of the wearable imaging system on a porcine wound model to demonstrate its efficacy.^26, 27^

### 4.1 Ethics statement

All animal experiments were conducted under the protocol approved by the University of California Davis (UC Davis) Institutional Animal Care and Use Committee (IACUC). All methods were performed in accordance with the UC Davis IACUC guidelines and regulations. All animal studies are reported in accordance with ARRIVE guidelines.

### 4.2 Animal Selection and Acclimatization

In each experiment, a female Yorkshire-Landrace-Duroc mix farm pig, weighing approximately 70–80 lbs and aged 4–5 months, was used. The pigs were acclimatized to the vivarium for 10 days before the experiment, during which they underwent daily behavioral training and were gradually accustomed to wearing a harness. Identification and initial weights were recorded for each animal. The pigs were singly housed and trained using positive reinforcement, such as food treats (fresh fruit and yogurt), to encourage standing still during future post-surgery checks.

### 4.3 Pre-Surgical Procedure

Prior to surgery, the pig was fasted for 12 hours and then induced under general anesthesia. Endotracheal intubation was performed, and anesthesia was maintained using masked inhalation of Isoflurane (1–5%) in 100% oxygen. Vital signs, including heart rate, respiratory rate, and temperature, were monitored every 15 minutes during the procedure. A blood sample was collected at the start of the surgery. The surgical area was clipped, depilated, and prepped with 2% chlorhexidine scrubs, isopropyl alcohol rinses, and a final betadine spray.

Following preparation, eight circular, full-thickness excisional wounds, each with a diameter of 20 mm, were created on the dorsum of the pig, as shown in Figure 4a. These wounds were evenly distributed bilaterally, with four wounds on each side. The wounds extended through the subcutaneous layer to ensure full-thickness injury. Devices were applied to the first six wounds, starting from the head, while the two wounds closest to the tail served as physiological controls.

Standard-of-care wound dressings (Conformant 2 wound veil, Optifoam bolsters, and Tegaderm film dressings) were applied to the control wounds.

### 4.4 Device Setup and Post-Surgical Care

The imaging devices were attached to the wounds, and their operation was verified before discontinuing anesthesia. Figure 4b shows the experimental pig’s left side, where the system’s setup is demonstrated. A black pouch, attached to the pig’s harness, housed the controller and battery pack, which powered the imaging devices. The harness was placed over the pig’s shoulder, and two devices are visibly attached to the wounds on the back.

After the devices were secured and operational, the entire wound area was covered with cushion foam sheets and a compression dressing or pig jacket to protect the wounded region. Post-surgery pain management included administering buprenorphine at extubation and applying a fentanyl patch, which provided analgesia for 72 hours. The pig was monitored to ensure it could stand before being returned to the vivarium to start the experiment (Day 0). The total duration of each experiment varied depending on the specific objectives, however, devices will remain on the wounds for either 3 or 7 days, with battery packs replaced every 24 hours to maintain a continuous power supply.

### 4.5 Mid-Experiment Procedures

During the 7-day experiment, dressing changes were performed on Day 3 in the surgical room, where new devices were applied despite their potential operational lifespan exceeding this time. After dressing changes, the pig was returned to the vivarium, initiating a new experimental phase (Day 3).

### 4.6 Endpoint Procedures

At the end-point, the pig was briefly anesthetized to collect a blood sample before euthanasia using pentobarbital (*>* 100 mg/kg IV). Devices were retrieved, and sterile cotton swabs were used to collect bacterial samples from the wounds. The wound tissues were harvested and bisected for further analysis. Portions of the tissue were preserved in PBS for fresh tissue studies, fixed in 4% paraformaldehyde for histological analysis, or processed for RNA sequencing (RNA-seq) to evaluate gene expression profiles.

## 5 Conclusion

In this study, we successfully developed a wirelessly connected, wearable imaging system for continuous wound monitoring, designed to work with a bioelectronic actuator and machine learning algorithm for real-time, personalized wound treatment. The device design integrates a 64 MP sensor, plano-convex lens, and programmable illumination to enable image acquisition of 20 mm diameter wounds with a spatial resolution of 44.19 *µ*m. *Ex-vivo* and *in-vivo* experiments validated the system’s ability to visualize wound healing progression. The modular controller architecture supported multiple camera devices, facilitating simultaneous monitoring of several wounds. Integration with DeepMapper demonstrated successful automated analysis of wound healing stages and detection of complications such as infection during our 7-day experiments. With continued refinement and validation through clinical studies, this platform has the potential to revolutionize wound care practices by enabling remote assessment, early complication detection, and data-driven treatment optimization.

## 6 Acknowledgement

This research is supported by the Defense Advanced Research Projects Agency (DARPA) through Cooperative Agreement Number D20AC00003 awarded by the U.S. Department of the Interior (DOI), Interior Business Center.

## 8 Supplementary Materials

Figure S1 shows the iterations of the camera design. The optics are identical between all versions. The overall shape of the device and the electronic boards were improved during the project.

## Notes

### Competing Interest Statement

The authors have declared no competing interest.

